# Jointly modeling prevalence, sensitivity and specificity for optimal sample allocation

**DOI:** 10.1101/2020.05.23.112649

**Authors:** Daniel B. Larremore, Bailey K. Fosdick, Sam Zhang, Yonatan H. Grad

**Affiliations:** Department of Computer Science, University of Colorado Boulder, Boulder, CO, 80309, USA; BioFrontiers Institute, University of Colorado at Boulder, Boulder, CO, 80303, USA; Department of Statistics, Colorado State University, Fort Collins, CO, 80523, USA; Department of Applied Mathematics, University of Colorado Boulder, Boulder, CO, 80309, USA; Department of Immunology and Infectious Diseases, Harvard T.H. Chan School of Public Health, Boston, MA, 02115, USA

## Abstract

The design and interpretation of prevalence studies rely on point estimates of the performance characteristics of the diagnostic test used. When the test characteristics are not well defined and a limited number of tests are available, such as during an outbreak of a novel pathogen, tests can be used either for the field study itself or for additional validation to reduce uncertainty in the test characteristics. Because field data and validation data are based on finite samples, inferences drawn from these data carry uncertainty. In the absence of a framework to balance those uncertainties during study design, it is unclear how best to distribute tests to improve study estimates. Here, we address this gap by introducing a joint Bayesian model to simultaneously analyze lab validation and field survey data. In many scenarios, prevalence estimates can be most improved by apportioning additional effort towards validation rather than to the field. We show that a joint model provides superior estimation of prevalence, as well as sensitivity and specificity, compared with typical analyses that model lab and field data separately.

---

Prevalence is traditionally estimated by analyzing the outcomes from diagnostic tests given to a subset of the population. During analysis of these outcomes, the sensitivity and specificity of the test, as well as the number of samples in the survey, are incorporated into point estimates and uncertainty bounds for the true prevalence. In many cases, sensitivity and specificity are taken to be fixed characteristics of the test [1, 2]. However, sensitivity and specificity are themselves estimated from test outcomes in validation studies. As a result, they, too, carry statistical uncertainty, and that statistical uncertainty should be carried forward into estimates of prevalence [3, 4]. Since prevalence estimates may improve as sample size increases and with reduced uncertainty in the test characteristics, a fundamental study design question arises: given a fixed number of tests, how should one allocate them between the field and validation lab?

Here, we derive a Bayesian joint posterior distribution for prevalence and test sensitivity and specificity based on sampling models for both the field survey data and validation data. While others have demonstrated how to estimate prevalence from this model [4–6], we highlight the utility of this model for addressing the problem of how to allocate a fixed number of tests between the field and the lab to produce the best prevalence estimates. We demonstrate that, when the sensitivity and specificity of a test have not yet been well established, that the largest improvement in prevalence estimates could result from allocating samples to test validation rather than to the survey. Finally, we showcase how this joint model can produce improved estimates of sensitivity and specificity compared to models based only on the lab data.

## METHODS

Our goal is to estimate population seroprevalence (*θ*), test sensitivity (*se*), and test specificity (*sp*) by learning from the field survey data *X* and the validation data *V*. The field survey data *X* contains the number of positive tests (*n*_+_) out of *N*_field_ samples. The validation data *V* contains the number of true positives (*tp*) resulting from *N*_pos_ positive control samples, providing information on test sensitivity, and the number of true negatives (*tn*) resulting from *N*_neg_ negative control samples, providing information about test specificity. We employ Bayes’ rule for estimation

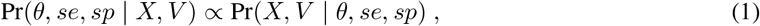

where we assume independent uniform priors on each of the parameters {*θ, se, sp*}. Survey data *X* and validation data *V* are collected independently via different processes but share the test’s sensitivity and specificity. We rewrite Eq. (1) as

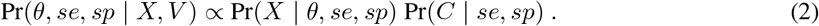

Given the parameter values {*θ, se, sp*}, the probability that a single random field test is positive is equal to the probability of obtaining a true positive or a false positive: *p* = *θse* + (1 – *θ*)(1 – *sp*). The number of positive outcomes after *N*_field_ independent tests is binomially distributed, so the probability of observing *n*_+_ positive tests is

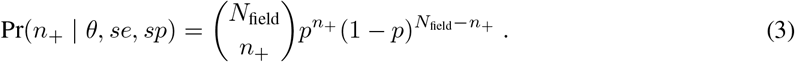

If the true test sensitivity is *se*, then the probability that a known positive sample produces a positive test outcome—a true positive—is *se*, while the probability of a false negative is 1 – *se*. The number of true positives *tp* in a set of *N*_pos_ independent positive validation test is also binomially distributed:

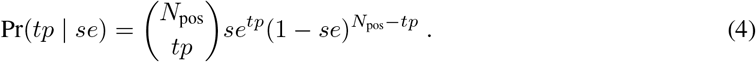

A parallel argument for true specificity *sp* and the outcomes of *N*_neg_ independent negative validation tests leads to the probability of observing *tn* true negatives:

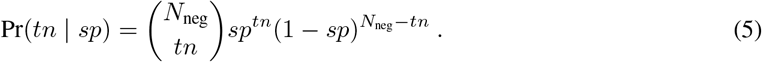

Substituting the probabilities in Eqs. (3), (4), and (5) into Eq. (2) and absorbing constants into the proportion, we obtain

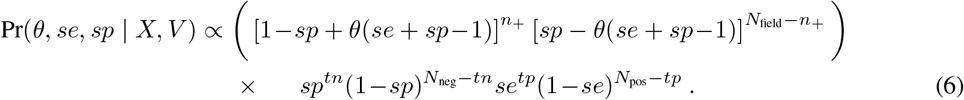

Equation (6) provides the form of the *joint* posterior distribution of *θ, se*, and *sp*, allowing one to learn simultaneously about these quantities and see how they depend on the data. Although the joint posterior distribution is not amenable to analytic computations (e.g. calculating expectations and variances), it is easily sampled using a Markov chain Monte Carlo (MCMC) algorithm. These posterior samples can then be used to estimate any summary statistics of interest, including point estimates (e.g., posterior means and modes) and credible intervals for the parameters. Posterior samples can further be passed as inputs into subsequent modeling tasks to account for uncertainty in prevalence [7, 8]. (See an An in-browser javascript calculator for computing the posterior distribution [9] and open-source code in R and Python [10].)

This model framework assumes that each diagnostic test is independent of the others and that the conditions in the field and validation are sufficiently similar that the diagnostic test has the same sensitivity and specificity in both. No attempts have been made to address or consider biases in the field sampling procedure itself.

## RESULTS

To demonstrate the impact of conducting additional validation tests, we computed the posterior distributions in Equation (6) for two scenarios in which field survey data consisting of 75 positive and 425 negative tests were analyzed using two sets of validation data. The first set was based on 100 validation tests and the other based on 200 validation tests, with tests split equally between positive and negative controls in both cases. Both validation data sets contained 94% true positives and 98% true negatives. The increase in validation samples resulted in a change in the 95% posterior credible interval for prevalence from [0.053, 0.182] to [0.082, 0.180] (Figure 1), corresponding to a 24% reduction in the credible interval width from additional validation data alone.

**FIG. 1.**
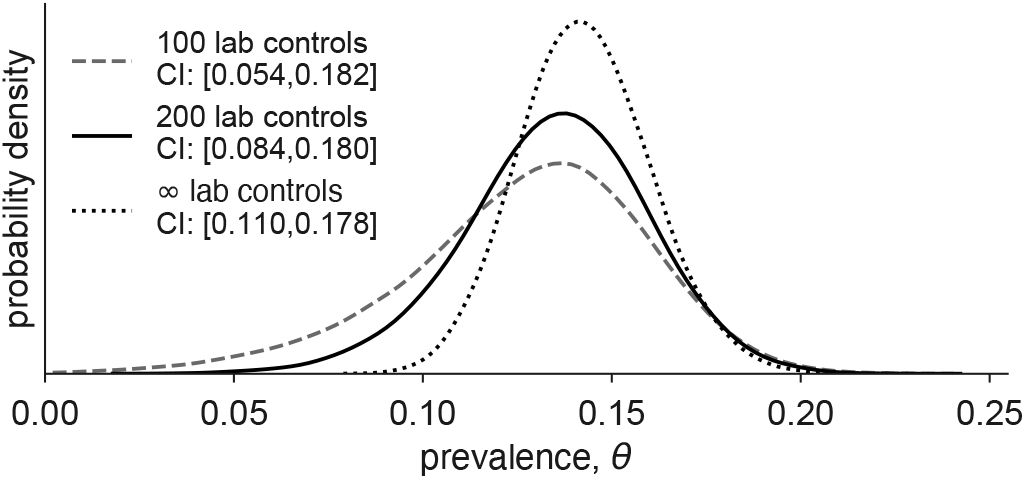
Increased validation effort decreases prevalence uncertainty. Prevalence estimates from 75 (*n*_+_) positives in 500 (*N*_field_) field samples, using validation outcomes of {*tp, tn*} = {47, 49} based on *N*_neg_ = *N*_pos_ = 50 samples (dashed line), {94, 98} based on *N*_neg_ = *N*_pos_ = 100 samples (solid line). Widths of 95% credible intervals decreased by 24% (prevalence), 32% (sensitivity), and 34% (specificity) due to increased validation efforts. Dotted line shows a Bayesian analysis of the same data using point estimates of 94% sensitivity and 98% specificity, equivalent to infinite lab validation data, for reference.

To compare the results of finite validation efforts to the theoretical optimum of infinite validation tests, we computed a posterior distribution for prevalence using point estimates of 94% sensitivity and 98% specificity. This results in a decrease in the width of the posterior credible interval by an additional 30% (Figure 1). The marginal impact of each additional validation test on posterior prevalence uncertainty decreases as this theoretical limit is approached.

When there is a limit on the number of tests that a prevalence study can use, due to budget, time, throughput, or other constraints, it may be tempting to deploy as many tests as possible to the field. This follows an intuition that additional field samples will decrease uncertainty in estimates of *θ*. However, while that intuition is correct, additional validation samples will also indirectly decrease uncertainty in *θ* by reducing uncertainty around sensitivity and/or specificity. By taking posterior uncertainty as the quantity to be minimized, we can search over combinations of *N*_field_, *N*_neg_, and *N*_pos_, representing the numbers of field, negative control, and positive control tests, respectively. When the total number of tests *N* = *N*_field_ + *N*_neg_ + *N*_pos_ is fixed, only two sample sizes can be specified freely, which means that this sample allocation problem becomes a minimization over a two-dimensional grid.

To demonstrate the use of this approach, we considered the allocation of *N* = 1000 tests in a setting where sensitivity and specificity are suspected to be around se = 0.93 and *sp* = 0.98 (based on, for instance, a similar test constructed by the same manufacturer) and in a population with suspected prevalence of 0.15. We allocated *N*_pos_ and *N*_neg_ to positive and negative controls, respectively, with the remainder allocated to *N*_field_. We then sampled from the posterior distribution for *θ* in Eq. (6) conditional on data equal to the expected counts of *tp, tn*, and *n*_+_. From these posterior samples, we computed the width of the 90% credible interval, and recorded it, before continuing to a new choice of sample allocation. Through this process, we found that at least twice as many samples should be allocated to specificity validation (*N*_neg_) as compared to sensitivity validation (*N*_pos_), and that around 1/3 of the 1000 total tests should be used for validation instead of for the field study (Figure 2). These specific allocation recommendations do not generalize to other *N*, prevalence, or test characteristics, but the search procedure itself is fully generalizable.

**FIG. 2.**
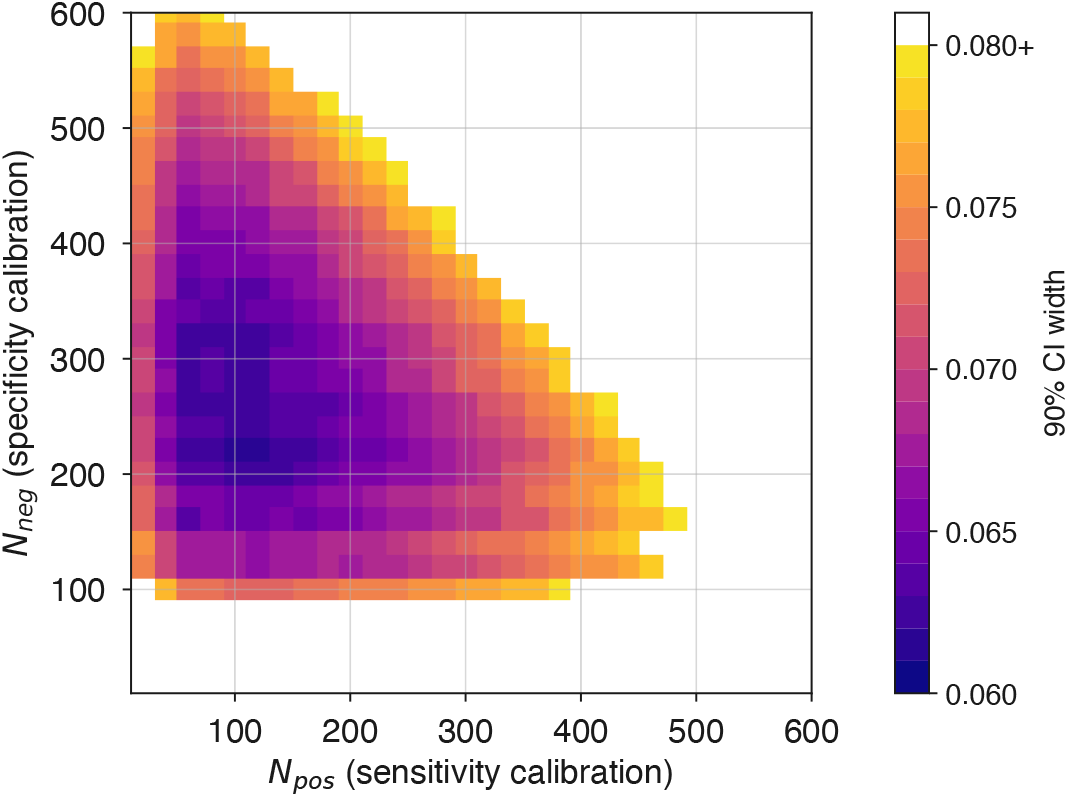
Optimized allocation of tests. Uncertainty in prevalence estimates, represented as 95% credible interval width, is shown as a heatmap for various allocations of *N* = 1000 tests, when prevalence is suspected to be 0.15, sensitivity 0.93, and specificity 0.98. Each pixel represents a choice of *N*_neg_ and *N*_pos_, where *N*_field_ = *N – N*_neg_ – *N*_pos_. Widths are indicated by color (see colorbar) with values larger than 0.09, or invalid choices of *N*_pos_ and *N*_neg_, in white. Each pixel was computed based on data equal to the expected test results for that allocation and using posterior samples from Eq. (6). Optimal allocations for the studied scenario favor allocation to negative controls over positive controls, with only 600-700 samples allocated to the field survey.

A second consequence of jointly modeling the validation and field data is that estimates of sensitivity and specificity may be affected by field survey data. Mathematically, this is because sensitivity and specificity appear in the probabilities of both the field and lab data sets in Eqs. (3), (4), and (5). To illustrate this point, we considered a scenario in which 95 of 100 negative controls were found to be negative during validation, resulting in a point estimate of specificity of 0.95, followed by a large study in a low prevalence area that resulted in only 10 positive tests out of 1000 samples. Such field data would appear inconsistent with the validation data, because even if prevalence were zero, one would expect 50 positives from 1000 field tests. However, an analysis based on Eq. (6) resolves this apparent inconsistency by inferring that the test’s specificity is likely to be higher than 0.95, with a posterior mean of 0.961 and posterior mode of 0.977 (Figure 3, solid line). For comparison, we also analyzed the validation data separately using a uniform prior on specificity, which produced a beta posterior distribution with a posterior mean of 0.941 and a posterior mode at 0.95 (Figure 3, dashed line).

**FIG. 3.**
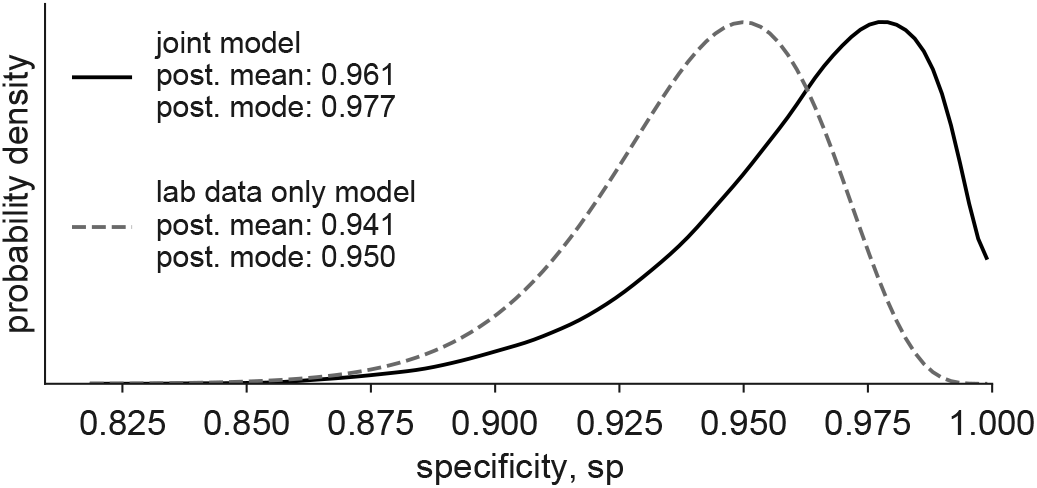
Test outcomes from the field affect estimates of sensitivity and specificity. Specificity estimates are shown for validation outcomes {*tp, tn*} = {100, 95} based on *N*_pos_ = *N*_neg_ = 100 controls analyzed independently of field data (dashed line; Beta posterior distribution) or jointly with *n*_+_ = 10 positives in *N*_field_ = 1000 field samples (solid line). While Fig. 1 illustrates the influence of lab validation data on prevalence estimates, this figure illustrates the less intuitive influence of field survey data on specificity estimates. This effect of field data is strongest on specificity when prevalence is low, and strongest on sensitivity when prevalence is high.

## DISCUSSION

The sensitivity and specificity of a diagnostic test are inferred from a finite number of validations tests. As a consequence, sensitivity and specificity themselves carry uncertainty, which affects the statistical interpretation of prevalence surveys in the field. Studies that use only point estimates of test characteristics can dramatically underestimate uncertainty around prevalence (Figure 1). Here, we showed how this issue can be ameliorated by jointly modeling field data and validation data using standard Bayesian techniques. Bayesian frameworks such this one can be used even when no validation data is available [11–13], can easily incorporate prior information about prevalence, sensitivity, or specificity from, other pilot or validation studies, and can jointly model the application of multiple diagnostic tests with different performance characteristics simultaneously [11]. These methods also avoid the need to rely on asymptotic approximations [1] in the process of calculating confidence intervals.

The direct inclusion of validation tests in prevalence estimation not only allows uncertain sensitivity and specificity to affect prevalence estimates (Figures 1 and 2), but also allows field data to affect sensitivity and specificity estimates (Figure 3). This underscores the importance of reporting the raw outcomes validation tests. The outcomes of validation tests should be included directly in publications that analyze field data whenever possible, motivated by statistical and reproducibility requirements.

By highlighting the marginal value of additional validation effort, joint models like Eq. (6) expose the tradeoff between collecting validation and field data when tests are limited. This simulation-informed approach to sample allocation allows a finite number of samples to be maximally utilized via strategic study design [7, 14, 15].

## ACKNOWLEDGEMENTS

The work was supported by the Morris-Singer Fund for the Center for Communicable Disease Dynamics at the Harvard T.H. Chan School of Public Health [to YHG].

## References

[1] Antoine Flahault, Michel Cadilhac, and Guy Thomas. Sample size calculation should be performed for design accuracy in diagnostic test studies. Journal of clinical epidemiology, 58(8):859–862, 2005.

[2] J Reiczigel, J Földi, and L O’zsva’ri. Exact confidence limits for prevalence of a disease with an imperfect diagnostic test. Epidemiology & Infection, 138(11):1674–1678, 2010.

[3] Walter J Rogan and Beth Gladen. Estimating prevalence from the results of a screening test. American Journal of Epidemiology, 107(1):71–76, 1978.

[4] Silvia Stringhini, Ania Wisniak, Giovanni Piumatti, Andrew S Azman, Stephen A Lauer, Helene Baysson, David De Ridder, Dusan Petrovic, Stephanie Schrempft, Kailing Marcus, Isabelle Arm-Vernez, Sabine Yerly, Olivia Keiser, Samia Hurst, Klara Posfay-Barbe, Didier Trono, Didier Pittet, Laurent Getaz, Francois Chappuis, Isabella Eckerle, Nicolas Vuilleumier, Benjamin Meyer, Antoine Flahault, Laurent Kaiser, and Idris Guessous. Repeated seroprevalence of anti-SARS-CoV-2 IgG antibodies in a population-based sample from Geneva, Switzerland. medRxiv, 2020.

[5] Stephen T Bennett and Mark Steyvers. Estimating covid-19 antibody seroprevalence in santa clara county, california. a re-analysis of bendavid et al. medRxiv, 2020.

[6] Jerome Levesque and David W. Maybury. A note on COVID-19 seroprevalence studies: a meta-analysis using hierarchical modelling. medRxiv, 2020.

[7] Daniel B Larremore, Bailey K Fosdick, Kate M Bubar, Sam Zhang, Stephen M Kissler, C. Jessica E. Metcalf, Caroline Buckee, and Yonatan Grad. Estimating sars-cov-2 seroprevalence and epidemiological parameters with uncertainty from serological surveys. medRxiv, 2020.

[8] Stephen M Kissler, Christine Tedijanto, Edward Goldstein, Yonatan H Grad, and Marc Lipsitch. Projecting the transmission dynamics of SARS-CoV-2 through the post-pandemic period. Science, 2020.

[9] https://larremorelab.github.io/covid19testgroup.

[10] https://github.com/LarremoreLab/bayesian-joint-prev-se-sp.

[11] Lawrence Joseph, Theresa W Gyorkos, and Louis Coupal. Bayesian estimation of disease prevalence and the parameters of diagnostic tests in the absence of a gold standard. American journal of epidemiology, 141(3):263–272, 1995.

[12] AJ Branscum, IA Gardner, and WO Johnson. Estimation of diagnostic-test sensitivity and specificity through bayesian modeling. Preventive veterinary medicine, 68(2-4):145–163, 2005.

[13] Peter J Diggle. Estimating prevalence using an imperfect test. Epidemiology Research International, 2011, 2011.

[14] Niel Hens, Ziv Shkedy, Marc Aerts, Christel Faes, Pierre Van Damme, and Philippe Beutels. Modeling infectious disease parameters based on serological and social contact data: A modern statistical perspective, volume 63. Springer Science & Business Media, 2012.

[15] Stephanie Blaizot, Sereina A Herzog, Steven Abrams, Heidi Theeten, Amber Litzroth, and Niel Hens. Sample size calculation for estimating key epidemiological parameters using serological data and mathematical modelling. BMC Medical Research Methodology, 19(51), 2019.

